# Overarching principles and dimensions of the functional organisation in the inferior parietal cortex

**DOI:** 10.1101/654178

**Authors:** Gina F. Humphreys, Rebecca L. Jackson, Matthew A. Lambon Ralph

## Abstract

The parietal cortex (PC) is implicated in a confusing myriad of different cognitive processes/tasks. Consequently, understanding the nature and organisation of the core underlying neurocomputations is challenging. According to the Parietal Unified Connectivity-biased Computation (PUCC) model two properties underpin PC function and organisation. Firstly, PC is a multi-domain, context-dependent buffer of time-and space-varying input, the function of which, over time, becomes sensitive to the statistical temporal/spatial structure of events. Secondly, over and above this core buffering computation, differences in long-range connectivity will generate graded variations in task engagement across subregions. The current study tested these hypotheses using a group ICA technique with two independent fMRI datasets (task and resting state data). Three functional organisational principles were revealed. Factor 1: inferior PC was sensitive to the statistical structure of sequences for all stimulus types (pictures, sentences, numbers). Factor 2: a dorsal-ventral variation in generally task-positive vs. task-negative (variable) engagement. Factor 3: An anterior-posterior dimension in inferior PC reflecting different engagement in verbal vs. visual tasks, respectively. Together the data suggest that the core neurocomputation implemented by PC is common across domains, with graded task engagement across regions reflecting variations in the connectivity of task-specific networks that interact with parietal cortex.

## Introduction

A long-history of neuropsychology and functional neuroimaging has implicated the parietal lobe in a confusing myriad of different cognitive processes and tasks. There is currently little clarity about the underlying core parietal neurocomputations. In a recent large-scale meta-analysis we investigated the functional organisation of the inferior parietal cortex (IPC) across multiple cognitive domains (Humphreys and Lambon Ralph 2015), revealing dorsal-ventral and anterior-posterior organisational graded variations in the types of task that engage IPC. Moreover, each subregion is engaged by multiple diverse tasks indicating that the region is not tessellated into distinct task-specific modules but rather the areas support domain-general computations that are called upon by different activities. Based on these results we proposed a unifying model of parietal function; the Parietal Unified Connectivity-biased Computation (PUCC). Here we test some of the central tenants of the model using two independent fMRI datasets as well as meta-analytic connectivity modelling.

There are three core assumptions of the PUCC model. The first proposes that the core local computation of the IPC supports online, multimodal buffering. Any time extended behaviour, whether verbal or nonverbal relating to internal or external cognition, requires some kind of internal representation of ‘the state of play’. Without a reliable representation of the current state it is impossible to check that that the state of the world has changed in the expected manner following the last action, to programme the next appropriate steps in the sequence towards the final goal, or to check that the state of the world has not changed dramatically in the interim such that a whole new goal needs to be instituted. Both automatized and executively-guided behaviours require access to an online buffered representation of the state of affairs. A second key notion relates to the possible broader computational differentiation across ventral (primarily temporal lobe) and dorsal (parietal) pathways. Specifically, the ventral processing routes generalise information across repeated episodes and input modalities, leading to context-independent representations. For example, in the case of semantic memory, multiple instances of a particular exemplar are generalised across time and contexts thereby allowing it to be recognised in highly variable situations and for information to be generalised across instances and contexts (Lambon Ralph et al. 2010; Buzsaki and Moser 2013; Lambon Ralph 2014). In contrast, the opposite is true for the parietal route, which appears to collapse information across items (i.e., statistically orthogonal to the ventral pathways) extracting item-independent time-, and space-varying structures (Buckner and Carroll 2007; Kravitz et al. 2011; Ueno et al. 2011; Bornkessel-Schlesewsky and Schlesewsky 2013). These two proposed features of the IPC – online buffering and extraction of item-independent time-/space-related statistics – can arise from the same computational process. For example, parallel distributed processing (PDP) models have demonstrated that through repeated buffering of sequential input the system becomes sensitive to the statistical structures of sequential information (McClelland et al. 1989; Botvinick and Plaut 2004, 2006; Ueno et al. 2011). In the action domain these statistical structures would support action schema, in the language domain it might result in the knowledge regarding phoneme or word order (depending on the time resolution over which statistics are computed) as well as number and spatial codings in other domains. A key prediction to be tested in this study was that in such models it is easier to process and buffer sequences that are typical of the statistical structure of the domain in question. Accordingly, we would expect activation in IPC to be (a) sensitive to violations in statistical knowledge and (b) to do so across multiple domains.

There are already hints from past studies that parietal cortex is sensitive to the temporal structure of events. For instance, IPC has been shown to respond when a word in a sentence is unexpected (Kuperberg et al. 2003; Hoenig and Scheef 2009), when ordering pictures into the correct sequence (Tinaz et al. 2006; Melrose et al. 2008; Tinaz et al. 2008), to scrambled motor sequences compared to learned sequences (Gheysen et al. 2010), or to violations in an expected visual sequence (Bubic et al. 2009). Furthermore, the notion that parietal cortex buffers context-dependent information is in accordance with several more domain-specific theories. For instance, IPC has been proposed as an “episodic buffer” of multi-modal episodic information (Wagner et al. 2005; Vilberg and Rugg 2008; Shimamura 2011), and others suggest that IPC acts as a phonological buffer/sensory-motor interface for speech (Baddeley 2003; Hickok and Poeppel 2007; Rauschecker and Scott 2009). Whilst domain-specific theories have been useful to account for findings from that domain of interest, they fail to explain how and why disparate cognitive domains coalesce in IPC subregions and thus what types of domain-general neurocomputations underlie processing across tasks (Corbetta and Shulman 2002; Humphreys and Lambon Ralph 2015, 2017).

A third hypothesis in the PUCC model is that although there might be a common overarching parietal neurocomputation, different parietal sub-regions show variations in processing based on graded variations in long-range connectivity. Previous computational models have demonstrated that even when there is a common overarching computation across a layer of units, differences in long-range connectivity generate graded variations in emergent function (Plaut 2002). Such connectivity variations might explain differences in the locus of activation in task-based studies (Bzdok et al. 2013; Humphreys and Lambon Ralph 2015). For example, tasks involving tool-use have been shown to overlap with numerous other tasks in dorsal parietal cortex (top-down attention, executive semantics, phonology, numerical calculation), yet the centre of mass of this cluster spreads towards motor and somatosensory areas (Humphreys and Lambon Ralph 2015). Thus, although there may be a high degree of overlap across tasks, the spread of activation for each will vary depending on the task-specific networks that connect to parietal cortex.

Such connectivity variations across parietal sub-divisions have been demonstrated using structural and functional connectivity measures. Angular gyrus (AG), supramarginal gyrus (SMG) and intraparietal sulcus/superior parietal lobule (IPS/SPL) have been shown to engage partially distinct neural networks: the AG forms part of the default mode network, the SMG forms part of a cingulo-opercular system and IPS/SPL is part of a fronto-parietal control system (Vincent et al. 2008; Spreng et al. 2010; Uddin et al. 2010; Cloutman et al. 2013; Power and Petersen 2013). There is some evidence that the transition between regions in terms of their connectivity profile is graded, rather than sharp in nature (Daselaar et al. 2013). Such connectivity-driven variations in function might also explain differences found between anatomically-proximate subregions (Uddin *et al.* 2010; Caspers et al. 2011; Cloutman *et al.* 2013): dorsal AG has been found to show positive activation for tasks involving semantic decisions on words and pictures, whereas middle AG is deactivated by both tasks, and ventral AG is activated by pictures but not words (Seghier et al. 2010).

In the current study, three independent datasets and a combination of methods were used to investigate these three core assumptions of the PUCC model. The first method used task-based fMRI. If the parietal cortex is sensitive to the statistical regularity of sequential input then it should more easily process sequential information with a regular statistical structure compared to structures that violate statistical likelihood. To test this, sequences of items were presented with either a regular structure or one were the structure was violated. We predicted that PC should show stronger responses to violated sequences of information compared to the normal sequences. Also, the model assumes that processing within PC is largely domain general but there may be some task differences based on variations in connectivity. To test this claim, different types of sequences were presented; comprising words, pictures or numbers. We predicted that whilst the region as a whole should be sensitive to sequence violations across tasks there may be differences in the pattern of activation across tasks based on variations in the task-specific network that interact with parietal cortex. To test our predictions, the data were analysed using a group spatial ICA. ICA has the advantage of being a data-driven method which can separate signal from noise components associated with movement or physiological fluctuations. As a result ICA has been shown to possess increased sensitivity compared to standard GLM techniques (McKeown et al. 2003). An additional advantage is that ICA can distinguish between distinct components with partial spatial overlap based on variations in time-courses (Leech et al. 2011). This point is significant because if sub-divisions are graded we expect some degree of spatial overlap across sub-regions. Therefore, task-ICA was used to investigate the functional networks involved in processing sequence violations across domains. After establishing the presence of distinct functional PC networks using the task data, an independent resting state dataset was used to independently verify the results.

## Methods

### fMRI task data

#### Participants

Twenty-participants took part in the study (average age = 24.4, SD = 4.79; N female = 16). All participants were native English speakers with no history of neurological or psychiatric disorders and normal or corrected-to-normal vision.

#### Task design

The participants completed three experimental tasks (sentence task, picture task, and number task) in separate scan sessions, the order of which was counterbalanced across subjects. In each task, on a given trial, a sequence of items (words, pictures, or numbers) was visually presented one item at a time with either a familiar structure (normal sequences) or a violated structure in which the final item from each sequence was taken from a different item. The participants’ task was to determine if the sequence was coherent. The sequences were selected from the most accurate subset from a pilot experiment. An example trial from each task is shown in Figure 1.

**Figure 1.**
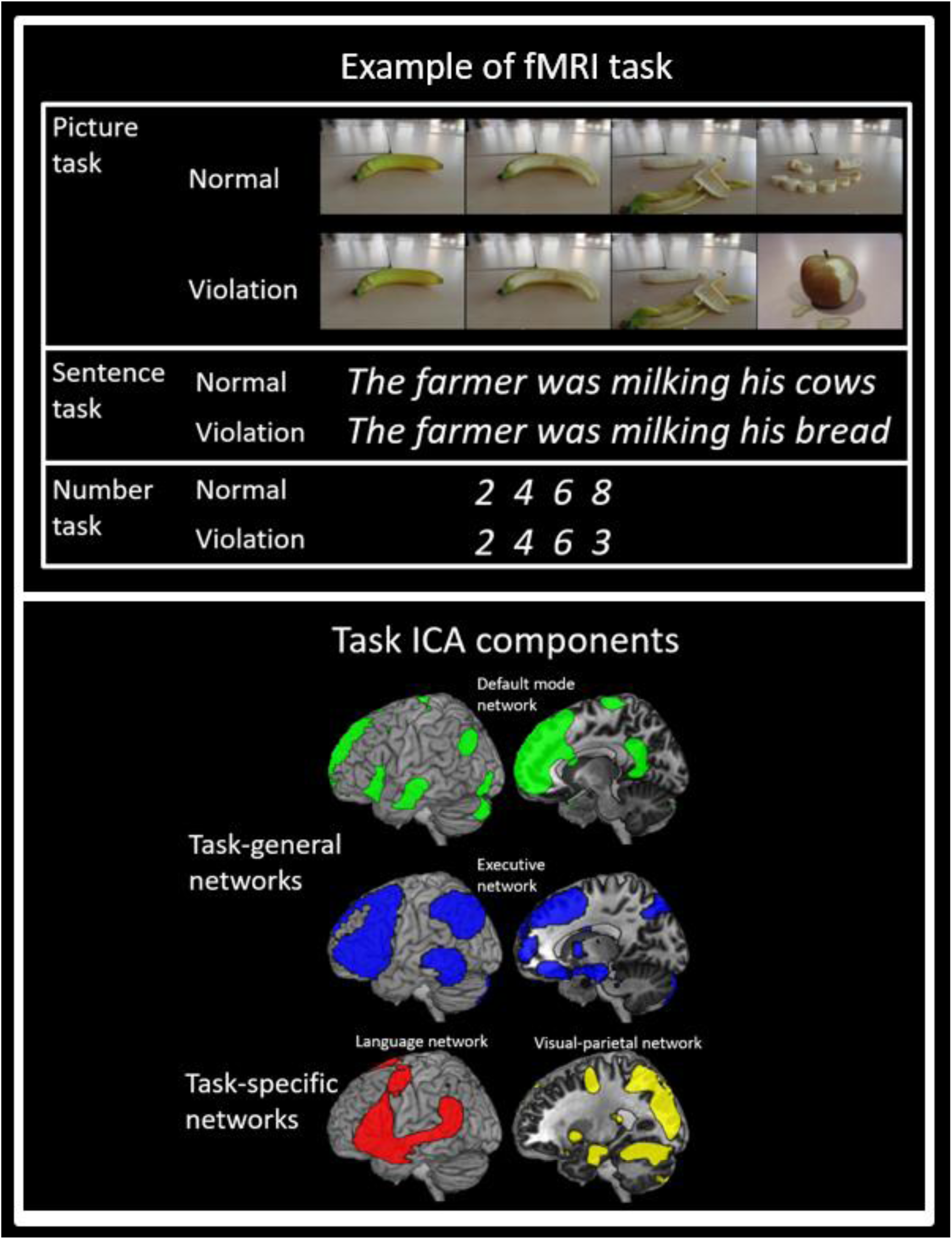
Top panel: An example from one trial for each of the tasks. Bottom panel: The task-general and task-specific ICA components (cluster corrected, p < .05).

#### Sentence task

to ensure a high degree of statistical regularity, sentences were selected in which the final word in the sentence had a high cloze probability and was thus highly predictable (e.g. *He loosened the tie around his <u>neck</u>*). The stimuli were a subset of the high cloze probability items included in Block and Baldwin (2010) (average cloze probability = 0.94, SD = 0.01). The sentence length varied from 6-10 words (average length = 8.4 words, SD = 1.0).

#### Picture task

a series of four colour pictures depicted the occurrence of real-life, everyday events with a clear causal structure, i.e., the events could not plausibly occur in a different order (e.g. a banana being peeled, a house being built, etc.). The images consisted of stills taken from freely available short online video clips downloaded from youtube.com. In each case, the event of interest was the central focus of the videos and there was minimal distracting background information.

#### Number task

a series of 4 numbers involving low-digit multiplication (e.g. *2 4 6 8*) or addition (e.g. *1 2 3 4*). Low-digit multiplication and addition have been shown to be automated skills, the solutions to which can be easily retrieved from memory (Simon et al. 2002; Dehaene et al. 2003). Note that the high accuracy scores for the task (see Results) confirm that the sequences were easily recognisable.

#### Task procedures

There were 42 items per condition presented using an event-related design with the most efficient ordering of events determined using Optseq (http://www.freesurfer. net/optseq). Null time was intermixed between trials and varied between 2 and 18 seconds (average = 4.59 seconds, SD = 3.06) during which a fixation cross was presented. For the picture and number task each of the four items in the sequence was presented for 900ms (total length = 3.6 seconds). The word-sequences in the sentence task contained between 6 and 10 words presented at a rate of one word every 360ms such that the maximum trial duration matched the picture and number task. Every item was followed by a “?” for 1.4 seconds at which point the participants provided a YES/NO button response.

#### Task acquisition parameters

Images were acquired using a 3T Philips Achieva scanner using a dual gradient-echo sequence, which is known to have improved signal relative to conventional techniques, especially in areas associated with signal loss (Halai et al. 2014). 31 axial slices were collected using a TR = 2.8 seconds, TE = 12 and 35ms, flip angle = 95°, 80 × 79 matrix, with resolution 3 × 3 mm, slice thickness 4mm. Across all tasks, 918 volumes were acquired in total, collected in six runs of 428.4 seconds each. B0 images were also acquired to correct for image distortion.

### Task data analysis

#### Preprocessing

The dual-echo images were first B0 corrected and then averaged. Data were analysed using SPM8. After motion-correction images were co-registered to the participants T1. Spatial normalisation into MNI space was computed using DARTEL (Ashburner 2007), and the functional images were resampled to a 3 × 3 x 3mm voxel size and smoothed with an 8mm FWHM Gaussian kernel.

#### General Linear Modelling

The data were filtered using a high-pass filter with a cut-off of 190s and then analysed using a general linear model. At the individual subject level, each condition for each task was modelled with a separate regressor (normal, violated) with time and dispersion derivatives added, and events were convolved with the canonical hemodynamic response function. Each sequence was modelled as a single event. Motion parameters were entered into the model as covariates of no interest. To investigate the effect of violation the contrast of violation sequences > normal sequences was computed in a whole-brain analysis (uncorrected, p < .001), with a significant cluster extent estimated using Alphasim with α < .05 and a brain mask applied. More targeted analyses were also conducted using the parameter estimates.

#### Task group spatial ICA

The pre-processed fMRI data were analysed in a group spatial ICA using the GIFT toolbox (http://mialab.mrn.org/software/gift) (Calhoun et al. 2001) to decompose the data into its components, separately for each task. GIFT was used to concatenate the subjects’ data and reduce the aggregated data set to the estimated number of dimensions using PCA, followed by an ICA analysis using the infomax algorithm (Bell and Sejnowski 1995). There were found to be 9 non-noise components for the number task, 11 for the picture task, and 13 for the sentence task. One sample t-tests were used to identify areas that significantly contributed to each component (cluster corrected, p < .05). The thresholded t-maps were then inspected and verbal labels were assigned to each network. Where possible labels were used which were consistent with those used frequently elsewhere in the literature (e.g. default-mode network (DMN), motor network, visual network, language network, saliency network) (Power et al. 2011; Lee et al. 2012; Yeo et al. 2013).

Certain components were found to be common to all tasks (see Supplementary Figure 1), we shall therefore refer to these as task-general networks. These closely resemble those that are commonly labelled as a DMN component and a fronto-parietal executive control component (described in detail in the Results). An additional network resembling that commonly referred to as the saliency network was also present, however this component was found to be insensitive to any task-manipulation and was therefore not included in further analyses. In addition to the task-general components we also identified two task-specific left parietal components (i.e., components which were not common to each task); a network that we have labelled as the “language component” from the sentence task, and a “visual-parietal component” from the picture task (described in detail in the Results).

In order to interrogate the cognitive signature of each component 12mm spheres were defined around the peak coordinates from all components of interest and these were used as ROIs to test for significant effects of conditions. Finally, we also examined how parietal networks might interact with one another or with other networks in the brain (e.g. visual or auditory) by performing a cross-correlation analysis of the average time-series for these components (parietal or non-parietal).

### Resting state data

#### Participants, procedures and acquisition parameters

Seventy-eight participants completed the resting state scan (average age = 25.23, SD = 5.55; N females = 57). During the scan the participants were instructed to keep their eyes open and look at the fixation cross. The data acquisition parameters for the resting state scan were identical to the experimental task. The scan consisted of a single 364 second scan session of 130 volumes.

#### Data Analysis

##### Preprocessing

Preprocessing was performed using SPM8 and the Data Processing Assistant for Resting State fMRI (DPARSF Advanced Edition, V2.3) toolbox (Chao-Gan and Yu-Feng 2010). Compared to the task data, additional preprocessing steps were carried out on the resting state data to minimise the influence of distance-dependent increases in correlations due to motion, which are considered problematic in resting state data. Thus, several procedures were adopted; censoring, global signal regression, 24 motion parameter regression and scrubbing of high motion time points. These methods have been shown to greatly reduce the effects of motion (Weissenbacher et al. 2009; Van Dijk et al. 2012; Yan et al. 2013; Power et al. 2014).

The images were first slice-time corrected, realigned and coregistered to the subjects T1 using SPM. Censoring was applied using a threshold of greater than 3mm of translation or 1 degree of rotation. This resulted in the exclusion of 6 participants from further analysis. Using DPARSF, images were normalised using DARTEL, smoothed with a 8mm full-width half maximum (FWHM) Gaussian kernel, and filtered at .01 - .08 Hz (Satterthwaite et al. 2013). Nuisance covariates were regressed out. These included covariates for 24 motion parameters, white matter, CSF and global tissue signal and also the performance of linear detrending. The 24 motion parameters were calculated from the 6 original motion parameters using Volterra expansion (Friston et al. 1996) and have been shown to improve motion correction compared to the 6 parameters alone (Yan *et al.* 2013; Power *et al.* 2014). Additional covariates were included for outlier time points with a with a z-score greater than 2.5 from the mean global power or more than 1mm translation as identified using the ARtifact detection Tools software package (ART; www.nitrc.org/projects/artifact_detect).

##### Resting state ICA

The goal of the resting state ICA analysis was to use an independent dataset to verify the AG functional subdivisions identified by the task ICA. The ICA was carried out on the preprocessed resting state data using the same method as the task data. This analysis identified 5 AG components the significance of which was tested using one-sample t-tests (FWE-corrected at the cluster level with a critical cluster level of .05). These 5 AG subdivisions were then used as ROIs for the task data to test for effects of violation and task.

## Results

### Behavioural results

Task performance was highly accurate across all experimental tasks (sentence task = 97%, SD = 3.3; picture task = 93%, SD = 5.8; number task = 91%, SD = 8.1). Nevertheless, there were some task differences: the sentence task was found to be significantly more accurate than the number task (t(19) = 3.17, p = .005), and marginally more accurate than the picture task (t(19) = 2.52, p = .02). The picture task was marginally more accurate than the number task when a Bonferroni correction was applied (t(19) = 2.17, p = .04).

In terms of reaction time, a 3 × 2 within-subjects ANOVA found a significant effect of task (F(38) = 18.13, p = .001), violation (F(38) = 7.71, p = .01), and a significant task x violation interaction (F(38) = 22.09, p = .001). Paired t-tests showed that responses to the sentence task were slower compared to the picture task (t(19) = 6.35, p = .001), and the number task (t(19) = 5.35, p = .001) which did not differ (t(19) = 1.06, p = .30). The interaction can be explained by an effect of violation in the picture task (t(19) = 5.75, p = .001), but no significant difference for the sentence task or number task (all ts < 2 ps > .05).

### GLM analysis

Compared to the normal sequences, the violation sequences elicited greater activation within IPC for all tasks (see Supplementary Figure 2 and Table 1), with overlap in medial posterior AG (PGp). There was also some overlap in the DLPFC, although this cluster did not survive the cluster correction for the number task. There were nevertheless some task differences. The violation effect was found to be significantly larger for the sentence task compared to the other two tasks combined (Sentences > Pictures + Numbers) in the anterior AG (PGa), left lateral frontal areas (inferior frontal gyrus and precentral gyrus), and right superior temporal gyrus. The left posterior middle temporal gyrus was also more strongly recruited for the sentence task, however this did not survive the cluster correction. The violation effect for the picture task was found to be greater than the sentence and number tasks combined (Pictures > Sentences + Numbers) in a network of bilateral visual areas (fusiform gyrus and visual cortex). There were no regions more responsive to the number task compared to the other two tasks. Therefore, these analyses support the hypothesis that IPC, especially AG, is sensitive to sequence violations but also suggests that there may be task-differences in the full network recruited.

### Task ICA analysis

#### Task-general networks

Certain parietal components were found to be largely overlapping across tasks (task-general networks). These resemble a DMN component (bilateral posterior AG, precuneus (PCC), medial frontal, mid-middle temporal gyrus (MTG)), a fronto-parietal executive control component (referred to as the executive network from here on, including left lateral frontal, AG/IPS, pMTG, and posterior superior frontal gyrus). The DMN and executive control network overlapped in AG, however the peak for the executive control network was found to be more dorsal. Both networks also overlapped in inferior frontal gyrus (IFG) and superior frontal gyrus (SFG), although the peak activation was more dorsal for the executive control network (Figure 1).

There were two parietal components that were task-specific in nature (see Figure 1). First, there was a visual-parietal network that involved visual cortex, SPL and PGp, which was present in the picture task alone. Secondly, there was a component identified from the sentence task data that clearly resembled an what is often referred to as the language network (left IFG, MTG, anterior AG) (Vigneau et al. 2006).

##### Sensitivity of task-general and task-specific networks to violation

In order to examine the cognitive signatures of each identified component, spheres were defined around the peak coordinates from the task-general (DMN, executive network, and saliency network) and task-specific networks (language, visual-parietal, and IPS-number) and these were used as ROIs to test for effects of violation.

#### Task-general parietal ROIs

The AG ROI for the executive network was more dorsal (in PGa, here on referred to as dorsal PGa) compared to the DMN ROI, which was in central AG (in PGp) (here on referred to as mid-PGp). Both ROIs showed an overall effect of violation, which did not interact with task: within the DMN, mid-PGp showed a significant effect of violation (F(1,19) = 5.73, p =.03) but no effect of task (F(2,38) = 1.75, p = .19) and no task x violation interaction (F(2,38) = .01, p = .99). Within the executive network, the dorsal-PGa ROI showed a significant effect of violation (F(1,19) = 18.63, p = .001), a marginal effect of task (F(2,38) = 3.01, p = .06) but no task x violation interaction (F(2,38) = 1.96, p = .16). The effect of task reflected moderately stronger activity for the picture compared to number tasks, however, this did not survive a Bonferronni correction (t(19) = 2.23, p = .04).

Despite the AG components of the executive network and DMN showing similar task-general sensitivity to sequence violations, the two sub-regions exhibited opposing directions of activation relative to fixation; activation for the executive network was significantly greater than zero for each condition (one-sampled t-test, all ts > 3.49, ps < .002), whereas the DMN elicits significant negative activation for each condition (one-sampled t-test, all ts > −3.68, ps < .002, although the sentence-violation condition only trended after Bonferroni correction was applied (t(19) = −2.8, p = .01). This suggests that whilst both areas show a similar effect of violation, the underpinning function of each subdivision is likely to differ; perhaps reflecting the fact that the dorsal PGa is part of the task-positive executive network, whereas the mid-PGp is part of the task-negative DMN. These results from all regions are presented in Figure 2.

**Figure 2.**
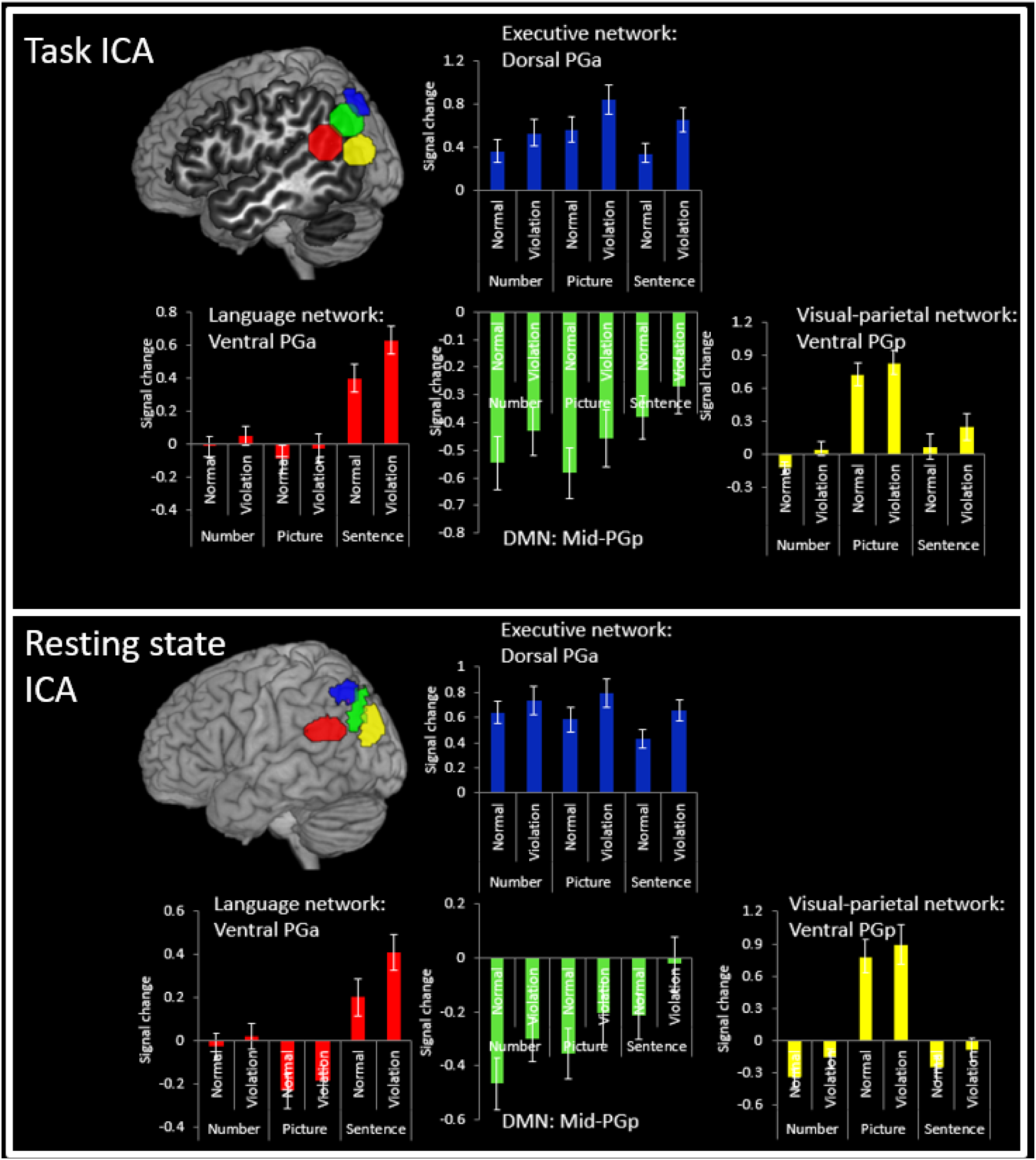
Percent signal change for the violation and normal sequences for each task within the Task ICA ROIs and resting-state ICA ROIs. The results show the same pattern for both methods.

#### Task-general non-parietal ROIs

Given that the AG component of the executive and DMN were both sensitive to sequence violations, further analyses were conducted on the non-parietal components of the networks so as to determine whether the effect was AG-specific or general to the whole of the network (see Figure 3).

**Figure 3.**
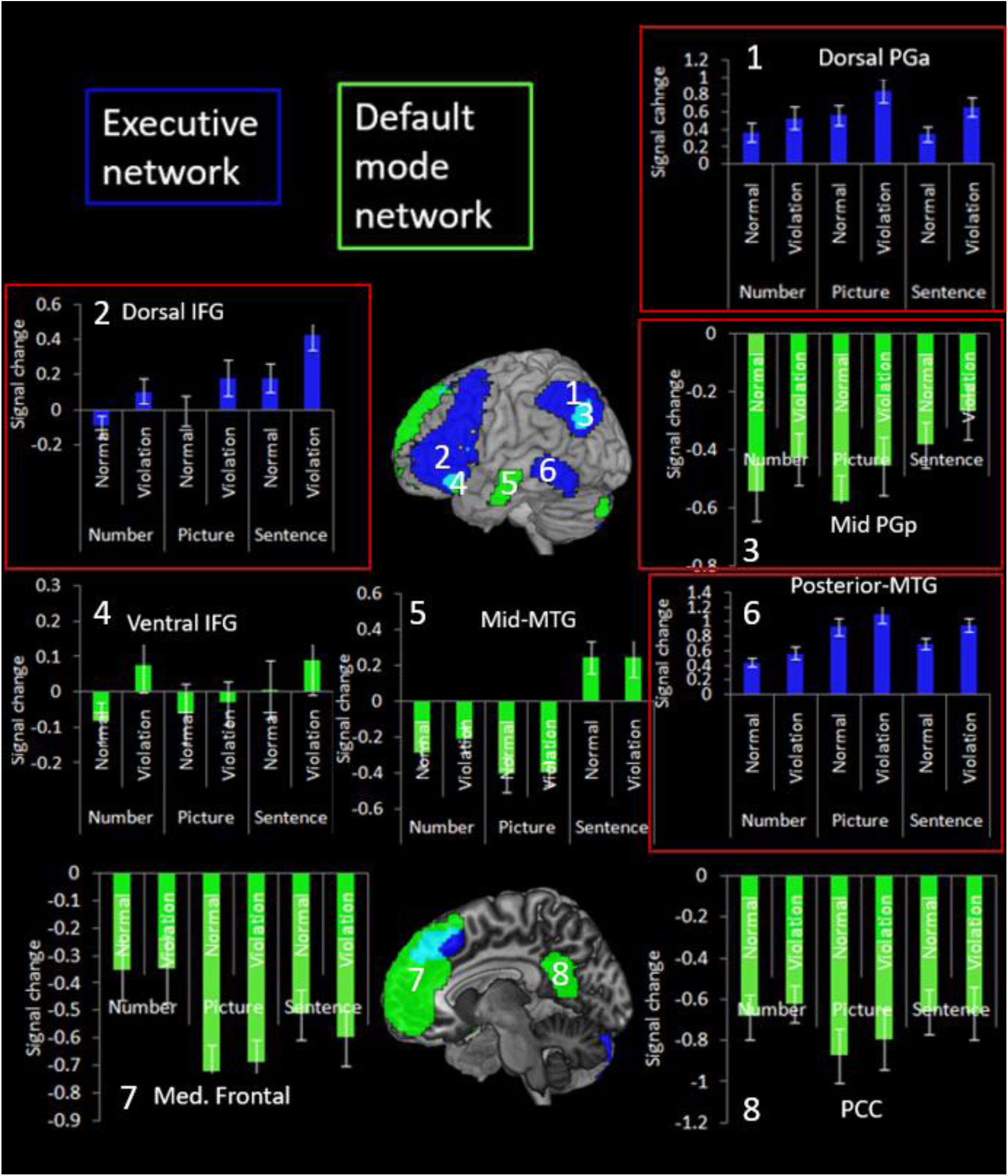
Percent signal change for the violation and normal sequences for each task within the executive network and default mode network. The regions to show a significant effect of violation are highlighted in red.

Within the DMN, no other region was found to be sensitive to violation. There were no significant effects for ventral IFG or PCC (all Fs < 2.58, ps > .13). Mid-MTG and medial frontal showed a significant effect of task but no effect of violation and no interaction (mid-MTG: task F(2,38) = 18.95, p = .001, condition F(1,19) = .30, p = .59, interaction F(2,38) = .47, p = .63; Medial frontal task F(2,38) = 5.35, p = .009, condition F(1,19) = .08, p = .78, interaction F(2,38) = .85, p = .44). For mid-MTG, sentences elicited greater activity compared to pictures (t(19) = 5.45, p = .001) and numbers (t(19) = 4.28, p = .001), which did not differ (t(19) = 1.58, p =.13). Within the medial frontal ROI, numbers elicited greater activity compared to pictures (t(19) = 3.52, p = .002). Therefore, the AG is the only DMN area to respond to sequence violations.

Unlike the DMN, all regions of the executive network showed task-general sensitivity to violation, with stronger activation for the violation condition compared to the normal sequences. Within the dorsal IFG, there was a significant effect of task (F(2,38) = 6.27, p =.004), violation (F(1,19) = 14.60, p = .001) but no significant task x violation interaction (F(2,38) = .20, p = .82). The task effect reflects greater activity for sentences compared to numbers (t(19) = 3.10, p = .006) and pictures (t(19) = 2.46, p = .02). Similarly, within posterior MTG there was a significant effect of task (F(2,38) = 18.11, p = .001) and violation (F(1,19) = 13.55, p = .002) and no significant task x violation interaction (F(2,38) = 2.54, p = .09). The task effect reflected reduced for numbers compared to pictures (t(19) = 5.74, p = .001) and sentences (t(19) = 4.56, p = .001).

#### Task-specific parietal ROIs

##### Language network

The anterior ventral AG (here on referred to as ventral PGa) showed a significant effect of task (F(2,38) = 26.92, p = .001), violation (F(1,19) = 10.75, p = .004), and a significant task x condition interaction (F(2,38) = 5.93, p = .006). The task effect reflects stronger activation for the sentence task compared to the picture task (t(19) = 6.15, p = .001) and the number task (t(19) = 7.32, p = .001). The interaction can be explained by a stronger effect of violation in the sentence task compared to the picture task (t(19) = 3.12, p = .006) and the number task (t(19) = 2.92, p = .009). One-sampled t-tests were used to examine whether the activation differed significantly from zero, and if so in which direction. This showed significantly positive activation for the sentence conditions only (ts > 4.72, ps < .001), with the picture and number conditions showing no difference from zero (ts < .92, ps > .37). Therefore, whilst this area is sensitive to violation overall, the effect is larger in the sentence task, which is also the only task to positively activate this region. This difference is likely explained by the fact that this region forms part of the language network and hence responds strongly to linguistic stimuli. These results are presented in Figure 2.

##### Visual-parietal network

The posterior ventral AG (here on referred to as ventral PGp) was specifically sensitive to the picture task. The results showed a significant effect of task (F(2,38) = 50.86, p = .001), violation (F(1,19) = 23.69, p = .001), but no task x violation interaction (F(2,38) = 1.16, p = .33). The picture task elicited stronger activation than the number task (t(19) = 10.02, p = .001) and the sentence task (t(19) = 7.62, p = .001), which differed marginally in favour of the sentence task (t(19) = 2.20, p = .04). Examinations of the direction of activation revealed significantly positive activation for the picture task only (ts > 6.35, ps < .001), with no modulation of the sentence and number tasks (ts < 2, ps > .05). Therefore, whilst this area is sensitive to violation overall, it shows a task-specific response to picture task likely due to the fact that this region is part of a visual processing network. These results are presented in Figure 2.

###### Cross-correlations

We examined how parietal networks might functionally relate to one another and also to the non-parietal neural networks by measuring the cross-correlations between each network’s time-series (Bonferroni corrected). Besides the networks mentioned above, we additionally included in this analysis the task-general auditory (bilateral auditory cortex) and visual networks (bilateral visual cortex) in order to have a common comparison across tasks and test whether each task differentially engaged each modality (e.g. picture tasks might correlate more strongly with visual components). Interestingly, like in the analyses described above, there was a dissociation in responses for the picture task compared to the sentence task (Figure 4. These showed a strong anti-correlation between the executive network (and related networks) and the DMN for both tasks, thus suggesting that activation of the executive network may lead to suppression in the DMN.

**Figure 4.**
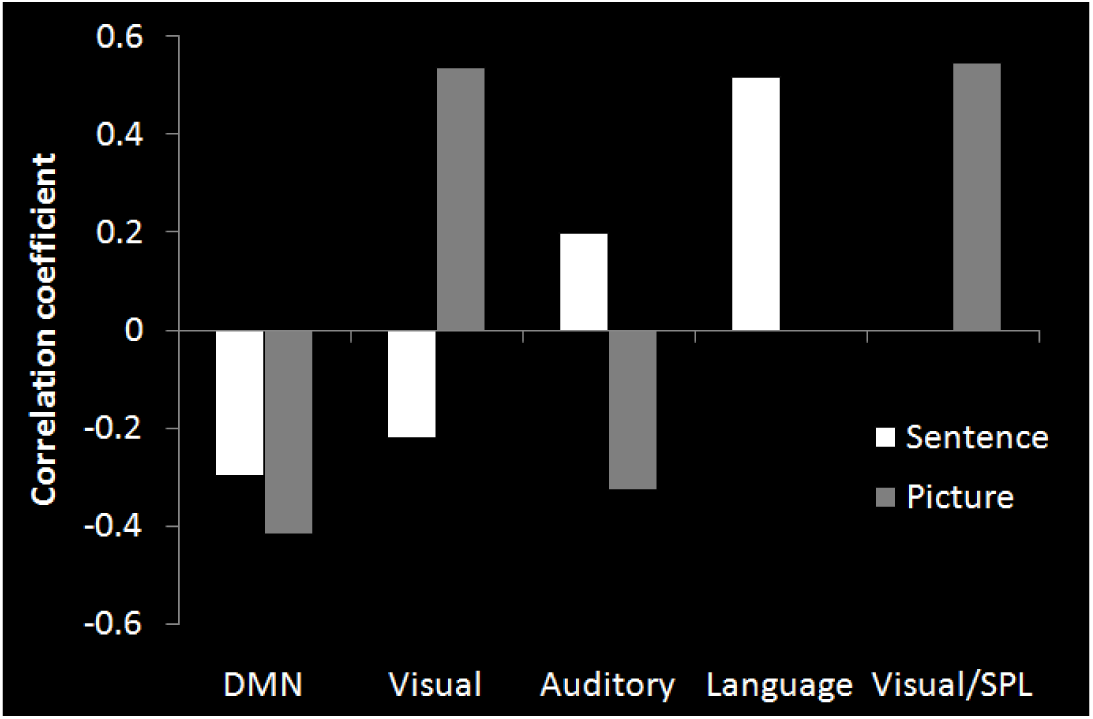
The cross-correlation with the average time-course of the executive network.

For the sentence tasks the executive network showed a significant positive correlation with the language network and also the auditory network (even though the sentences were visually presented), but a negative correlation with visual networks and DMN. Whereas, in the picture task the executive network instead correlated positively with visual networks and negatively with auditory networks and DMN. This suggests that whilst the executive network is task-general there is a shift in the networks that interact with it based on the varying demands of each task. Furthermore, when a network is not required for that particular task it becomes anti-correlated with the executive network.

###### Summary

The task ICA results suggest that IPC as a whole is sensitive to sequence violations. However, IPC subregions appear to be organised along dorsal-ventral and anterior posterior-dimensions. Specifically, dorsal PGa and mid-PGp showed task-general responses, yet the time-series of the two networks were anti-correlated and the sub-regions had opposing activation directions relative to rest: dorsal PGa was positively activated by all tasks whereas ventral mid-PGp was deactivated.

Whereas these areas showed a task-general response to violation, anterior and posterior portions of ventral IPC showed task-specific responses. Specifically, ventral PGa was only positively activated by sentence tasks, whereas ventral PGp responded positively to picture tasks alone. This pattern mirrors the variations in the networks that correlate with each sub-region. Specifically, ventral PGa is part of the language network and hence is positively activated for sentence tasks, whereas ventral PGp is part of a visual network and hence responds to picture tasks. There was also a dynamic, task-specific switching between the executive network and the other networks; for the sentence task the executive network correlated with language and auditory networks and was anti-correlated with visual areas, but for the picture tasks the opposite pattern was found.

##### Resting state ICA

The resting state ICA analysis was used to verify the presence of the functional sub-divisions using an independent dataset and in the absence of a task. This ICA revealed five components which involved IPC. Components 1 and 2 engaged partially overlapping regions of dorsal PGa. Component 1 included a similar network as the executive component from the task ICA analysis (lateral frontal, dorsal parietal, and pMTG), whilst component 2 was more restricted in size but still recruited lateral frontal and dorsal parietal areas. Component 3 engaged mid-PGp region and was similar to the DMN identified in the task-ICA analysis. Component 4 engaged ventral PGa and resembled the language network from the previous analysis. Finally, component 5 engaged ventral PGp and included a network of superior parietal and higher-level visual areas which included some of the same regions as the visual/SPL network from the task ICA analyses. Thus, there appeared to be strong correspondence between the networks identified in the task-based and resting state ICA analyses (see Figure 5).

**Figure 5.**
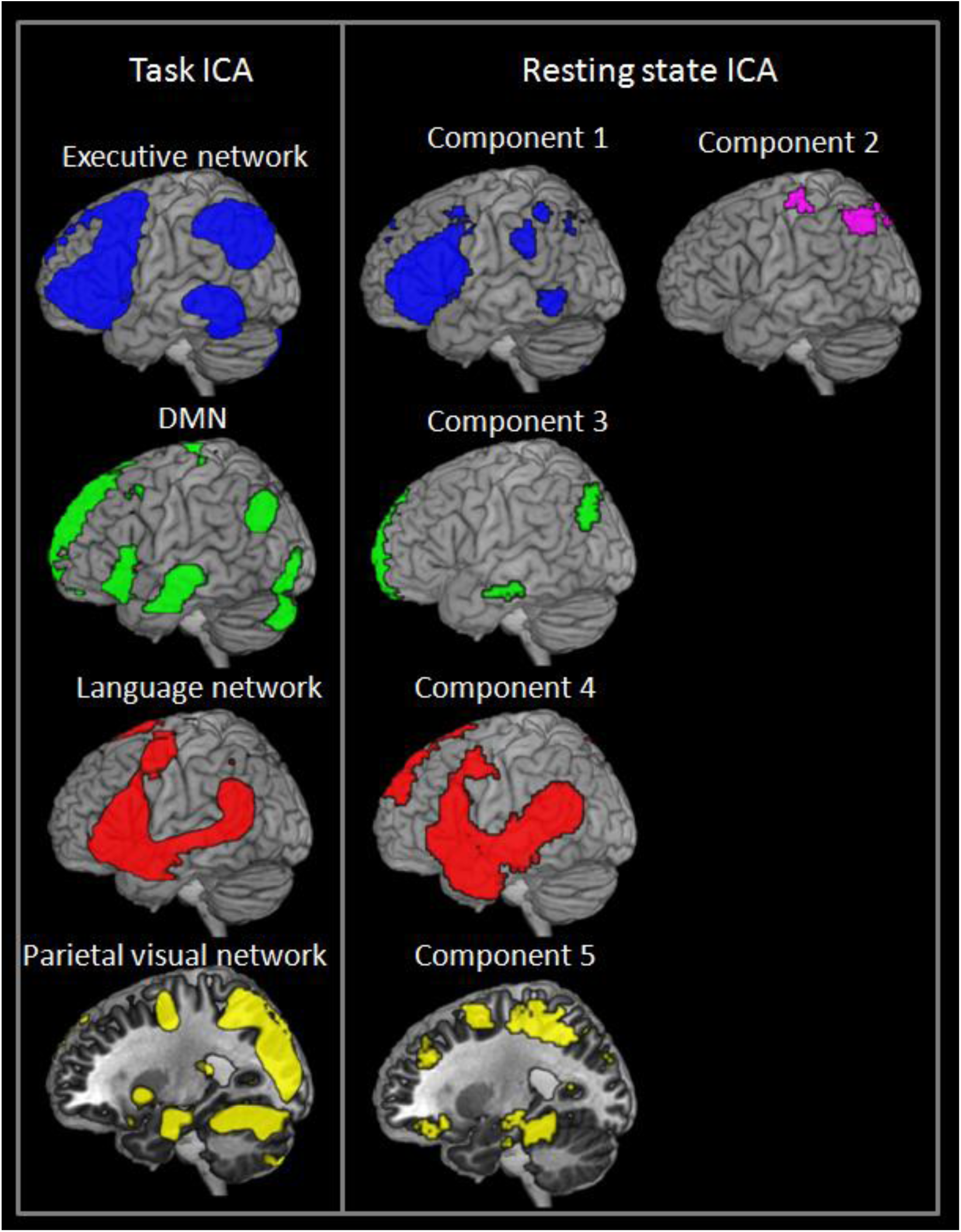
The correspondence between the task ICA and resting state ICA analyses.

The five rs-fMRI derived IPC sub-divisions were used as ROIs for the task-data to examine whether they showed a similar pattern of sensitivity to violation across tasks as those regions defined by the task-ICA (see Figure 2). Responses within components 1 and 2 were found to be similar to the dorsal PGa region from executive network in the task-ICA data. Both components 1 and 2 showed a significant effect of violation (Component 1: F(1,19) = 13.13, p= .002; Component 2: F(1,19) = 15.13, p = .001) but no effect of task (Component 1: F(2,38) = 1.57, p = .22; Component 2: F(2,38) = 1.36, p = .87) and no task x condition interaction (Component 1: F(2,38) = 1.98, p = .15; Component 2: F(2,38) = .94, p = .40). Activation was also found to be significantly positive compared to zero across all conditions (all ts > 3.81, ps < .002).

Responses within component 3 resembled those of mid-PGp DMN in the task data. There was a significant effect of condition (F(1,19) = 10.10, p = .005) but no effect of task (F(2,38) = 2.76, p = .08) and no task x condition interaction (F(2,38) = .142, p = .87). Responses tended to be negative relative to zero (all ts > 2.53, p < .01) except for the picture violation and sentence violation conditions which did not differ from zero (t < 1.81, p > .09).

The response within component 4 and 5 resembled the task-specific responses found for the language network and visual-parietal network from the task data, respectively. Specifically, for component 4 there was a significant effect of task (F(2,38) = 22.54, p = .001), violation (F(1,19) = 7.67, p = .01), and a significant task x violation interaction (F(2,38) = 4.86, p = .01). Paired t-tests showed greater activation for the sentence task compared to the number task (t(19) = 4.97, p = .001) and picture task (t(19) = 5.83, p = .001), and greater activation for the number task compared to the picture task (t(19) = 2.62, p = .02). Also, the effect of violation was significantly larger in the sentence task compared to the picture task (t(19) = 2.97, p =.008) or the number task (t(19) = 2.44, p = .02), although this was trending using a Bonferroni correction. Compared to zero, responses were significantly positive or trending for the sentence conditions (all ts > 2.34, p < .03), did not differ from zero for the number conditions (all ts <.34, p > .74), and were negative for the picture conditions (all ts > 2.52, p < .02).

For component 5 there was a significant effect of violation (F(1,19) = 19.83, p = .001) and task (F(2,38) = 57.46, p = .001) but no task x condition interaction (F(2,38) = .60, p = .55). Paired t-tests showed that the picture task elicited significantly greater activation compared to the sentence task (t(19) = 8.09, p = .001) and the number task (t(19) = 8.95, p = .001), which did not differ (t(19) = .93, p = .37). Against zero, activation was positive for the picture condition (all ts > 4.70, ps < .001), did not differ from zero for the sentence conditions (all ts < 2.1, p > .05), and was negative or trending for the number condition (all ts > 1.87, p < .08).

## Discussion

The multi-method approach used in this study revealed several key findings with regard the function of the inferior parietal cortex (IPC). Aligning with the predictions of the PUCC model (see Introduction), the highly convergent results can summarised in terms of three contributing factors.

### Factor 1: sensitivity to violation

multiple parts of IPC and the whole executive network are sensitive to violations in the structural coherence of sequences across domains; they respond more strongly to sequences in which the statistical regularity is violated compared to regular sequences. Despite this general property, the IPC has graded functional subdivisions as observed in the task ICA and replicated using the independently defined ROIs from the resting state ICA. Together these ROI analyses plot out two primary axes of IPC organisation (described next).

### Factor 2

A dorsal-ventral difference was established with more dorsal areas (dorsal PGa), forming part of the executive network and responding with positive activation to sequences in a domain general fashion. In contrast, more ventral areas (mid PGp) form part of the DMN and are deactivated by all tasks (though mid PGp was the only part of the DMN that was sensitive to sequential violation). Moreover, the executive network and DMN showed anti-correlated time-series. Together these results suggest that activation of the top-down executive network may lead to suppression of the DMN.

### Factor 3

the final factor to influence the results was an anterior-posterior dimension of organisation within ventral IPC. Ventral PGa formed a part of the language network and hence responded specifically to linguistic material (sentences), whereas ventral PGp was part of the visual/SPL network and hence responded to pictorial material alone. Also the language and visual-parietal networks selectively correlated with the executive network only when their preferred task was performed. This suggests task-dependent dynamic flexibility in the regions in their interaction with the core, multi-demand executive network.

The results show that IPC, together with the executive network, responded to task-general sequence violations. The PUCC model proposes that the IPC may form a neuroanatomically-graded multi-modal buffer thereby supporting a dynamic representation of the changing internal and external ‘state of affairs’. As a by-product of repeated events, this system will become sensitive to the temporal and spatial statistics (Plaut, 2003). Accordingly, sequences that violate statistical regularity are more effortful to process and thus elicit greater activation. The current data are consistent with existing studies finding that IPC responds to the statistical regularities of meaningful (words/picture sequences) and meaningless events (motor/visual sequences) (Kuperberg *et al.* 2003; Tinaz *et al.* 2006; Melrose *et al.* 2008; Tinaz *et al.* 2008; Bubic *et al.* 2009; Hoenig and Scheef 2009; Gheysen *et al.* 2010). Indeed, there is a growing body of evidence that IPC forms part of a context-related processing network. For instance, it responds more strongly to images with strong rather than weak contextual associations (Bar et al. 2008), or when subjects remember contextual associates of an item (Fornito et al. 2012), and is sensitive to event occurrence frequency (d’Acremont et al. 2013).

IPC responses were found to be task-general with some variations around the anterior and posterior edges. This supports the notion that there is a core underling IPC neurocomputation which is common across tasks (Walsh 2003; Cabeza et al. 2012; Humphreys and Lambon Ralph 2015) and argues against a highly “fractionated” or modular pattern of organisation (Nelson et al. 2012). Indeed, the current data appear inconsistent with any domain-specific theories of IPC function which, for example, suggest specialisation for semantic memory (Geschwind 1972; Binder et al. 2009), episodic memory (Wagner *et al.* 2005; Vilberg and Rugg 2008; Shimamura 2011), or numerical processing (Arsalidou and Taylor 2011). For example, it is hard to reconcile the notion that the AG is a store for semantic representations or alternatively the locus of magnitude processing, with the finding of equal deactivation for numerical and semantic tasks. Similarly, if the area was specialised for episodic information it is unclear why it would be sensitive to sequence violations which do not involve the retrieval of information relating to a specific episode. Together with previous cross-domain explorations of IPC function (Cabeza *et al.* 2012; Humphreys et al. 2015; Humphreys and Lambon Ralph 2015, 2017), the current data are more consistent with the notion of a domain-general process but with graded differences in function based on variations in connectivity to different AG sub-regions.

The ventral parietal cortex is involved in bottom-up/stimulus-driven and automatic task components (Cabeza *et al.* 2012; Humphreys and Lambon Ralph 2015). For instance, AG shows stronger activation for faster reaction times (Hahn et al. 2007) and is sensitive to a range of tasks with more automated tasks compared to executively demanding tasks, e.g. numerical fact retrieval vs. numerical calculation, or making semantic decisions on concrete vs. abstract words (Humphreys and Lambon Ralph 2015). In contrast, the executive network including dorsal IPC subregions are known to be involved in top-down processing, responding more strongly to difficult decisions or task demands across diverse domains and task-types (Fedorenko et al. 2013; Noonan et al. 2013; Humphreys and Lambon Ralph 2017). The relationship between the bottom-up and top-down networks is unclear but it is possible that when currently-buffered information cannot be automatically processed by the ventral IPC subregions and their connected networks (as in the case of sequence violations), then this triggers the involvement of top-down executive processing systems (see Humphreys & Lambon Ralph, 2015 for further discussion). Indeed, this is akin to the notion of a “circuit-breaker” proposed by Corbetta and Shulman (2002) in which the stimulus-driven network acts as an alerting system for top-down processing.

Two anatomical gradients of organisation were identified within IPC; dorsal-ventral and anterior-posterior. The fact that dorsal (IPS/SPL) and ventral parietal (AG/SMG) areas are functionally dissociable has been recognised by several models of parietal function (Corbetta and Shulman 2002; Cabeza et al. 2008; Humphreys and Lambon Ralph 2015). Indeed, dorsal and ventral parietal cortex connect with distinct cortical areas: AG forms part of the default mode network whereas IPS/SPL is part of a fronto-parietal control system (Vincent *et al.* 2008; Spreng *et al.* 2010; Uddin *et al.* 2010; Cloutman *et al.* 2013; Power and Petersen 2013). fMRI studies have also shown that dorsal IPC is associated with task-positive activation, whereas ventral IPC is typically associated with task negative-activation (Fox et al. 2005). The current findings demonstrate that rather than a sharp dissociation between dorsal (IPS/SPL) and ventral (AG/SMG) areas there is a graded shift in activation even within AG: regions towards IPS become positively activated and relate more strongly to the executive network compared to the DMN. The current results are consistent with a similar graded shift from negative to positive activation in AG observed for semantic tasks (Seghier *et al.* 2010), though the current study shows that this pattern is not specific to semantic tasks but rather a task-general feature.

The results also showed that the executive network and DMN had anti-correlated time-series. Likewise, resting-state studies have frequently shown that these networks are anti-correlated (Fox *et al.* 2005; Hampson et al. 2010), nevertheless this is some of the first evidence to show this dynamic interplay during task performance. Future studies are needed to answer the subsequent questions that arise from this repeated observation (see also Humphreys & Lambon Ralph, 2015). First, why are any brain regions deactivated at all? Two important possibilities include the observation that “rest” is not a neutral condition but rather allows in-scanner spontaneous cognition and internal processes and thus “deactivation” might reflect the fact that some active fMRI tasks do not share these cognitive processes (Buckner and Carroll 2007; Raichle and Snyder 2007; Binder *et al.* 2009; Andrews-Hanna 2012). Another possibility relates to the fact that regions tuned to task-irrelevant functions might be deactivated to save metabolic energy (Attwell and Laughlin 2001; Humphreys *et al.* 2015). This second possibility is consistent with the results found here for the anterio-posterior changes in function across the ventral IPC (and other findings: see Humphreys et al 2017). Ventral PGa is tuned more towards language whilst ventral PGp for visual tasks. When the active tasks matches their function then these regions exhibit positive activation whereas during other types of tasks they actually deactive.

A second puzzle is why the executive and DMN are often (though not always) anti-correlated, with the degree of DMN deactivation and executive network activation both correlated with task/item difficulty, regardless of task (Fedorenko *et al.* 2013; Humphreys and Lambon Ralph 2017). The PUCC model suggests that the two networks are often counterpointed because ventral IPC buffering for automatic activities, by definition, do not require working memory or ‘problem-solving’ mechanisms; whereas when an ongoing task becomes problematic the executive network is engaged and ongoing automatic buffering may be counterproductive for problem-solving and thus the buffering is temporarily suspended or suppressed. These notions are similar to previous suggestions for a ‘safety break’ mechanism formed through the dynamic interplay between IPS and IPC, and triggered when an unexpected event or stimulus is encountered in the ventral network (Corbetta and Shulman 2002).

The third question relates to what types of task generate task-positive activation in ventral IPC regions and by extension the DMN. These regions are most often associated with task-related deactivation and thus, understanding the conditions under which task positive responses are observed, might provide critical clues about these regions’ core function. This study and related investigations (e.g., Humphreys et al 2017) provide the first evidence for modality-related variations of processing within AG, which is frequently considered as a modality-general processing area (Binder and Desai 2011) and align with recent proposals that the DMN, more generally, is a multifaceted entity which fractionates depending on the nature of the task that is compared to “rest” (Buckner et al. 2008; Humphreys *et al.* 2015; Axelrod et al. 2017). The current study observed this type of fractionation along the ventral IPC region (see also Humphreys et al., 2017): ventral PGa exhibited deactivation in all conditions except for the language sequences when it was positively activated; ventral PGp showed exactly the reverse pattern. Such results run counter to any single cause or domain-general reason for deactivation but are consistent with notions that areas unnecessary for the current task are deactivated, perhaps to minimise cognitive interference and/or to save metabolic energy energy (Attwell and Laughlin 2001; Humphreys *et al.* 2015; Humphreys and Lambon Ralph 2015). The mid-AG remains something of a mystery in that it deactivated across all conditions (albeit being sensitive to sequence violations like the entire IPC region) and is one of the areas consistently associated with the DMN (Buckner et al 2008). Future cross-domain comparative fMRI studies are required to establish which subtypes of task generate positive activations in the mid-AG and whether these tasks are selective to this IPC subregion, as ventral PGa and PGp appear to be for language and visual tasks, respectively. Possibilities include mind-wandering or other forms of internally-directed cognition (Andrews-Hanna 2012), vivid episodic/autobiographical recall (Wagner *et al.* 2005; Vilberg and Rugg 2008), or future thinking (Buckner and Carroll 2007).

The final question to be considered here pertains to what drives these graded anterior-posterior and superior-ventral graded functional variations across the IPC region? The PUCC model, like other proposals (Cabeza *et al.* 2012) assumes that, whilst the IPC might have a core basic neurocomputation (e.g., buffering of current information), subregions come to exhibit gradedly different responses depending on their pattern of long-range connectivity. This computational principle has been demonstrated previously for PDP models of semantic representation (Plaut, 2003). In terms of the anterior-posterior AG gradient, ventral PGa responded positively to the sentence task presumably due to input from the verbally-related posterior temporal (STS/MTG) areas, whereas ventral PGp exhibited activation for the picture task perhaps reflecting greater connectivity to visually-related occipital/occipitoparietal regions (Ruschel et al. 2014). In a similar vein, the strong dorsal-ventral IPC variation is likely to reflect differential connectivity, with stronger connections from dorsal AG/IPS regions to DLPFC, thus forming the foundation for the multi-demand, executive network (Uddin *et al.* 2010; Yeo et al. 2011).

To conclude, the inferior parietal cortex exhibits cross-domain sensitivity to the structural coherence of sequential input, consistent with a multimodal buffering computation. This generalised function is conditioned across dorsal-ventral and anterior-posterior dimensions in keeping with variations in long-range connectivity.

## Acknowledgements

This research was supported by an MRC Programme grant to MALR (MR/R023883/1) and a British Academy fellowship to RLJ (pf170068).

## Supplementary

**Supplementary Figure 1.**
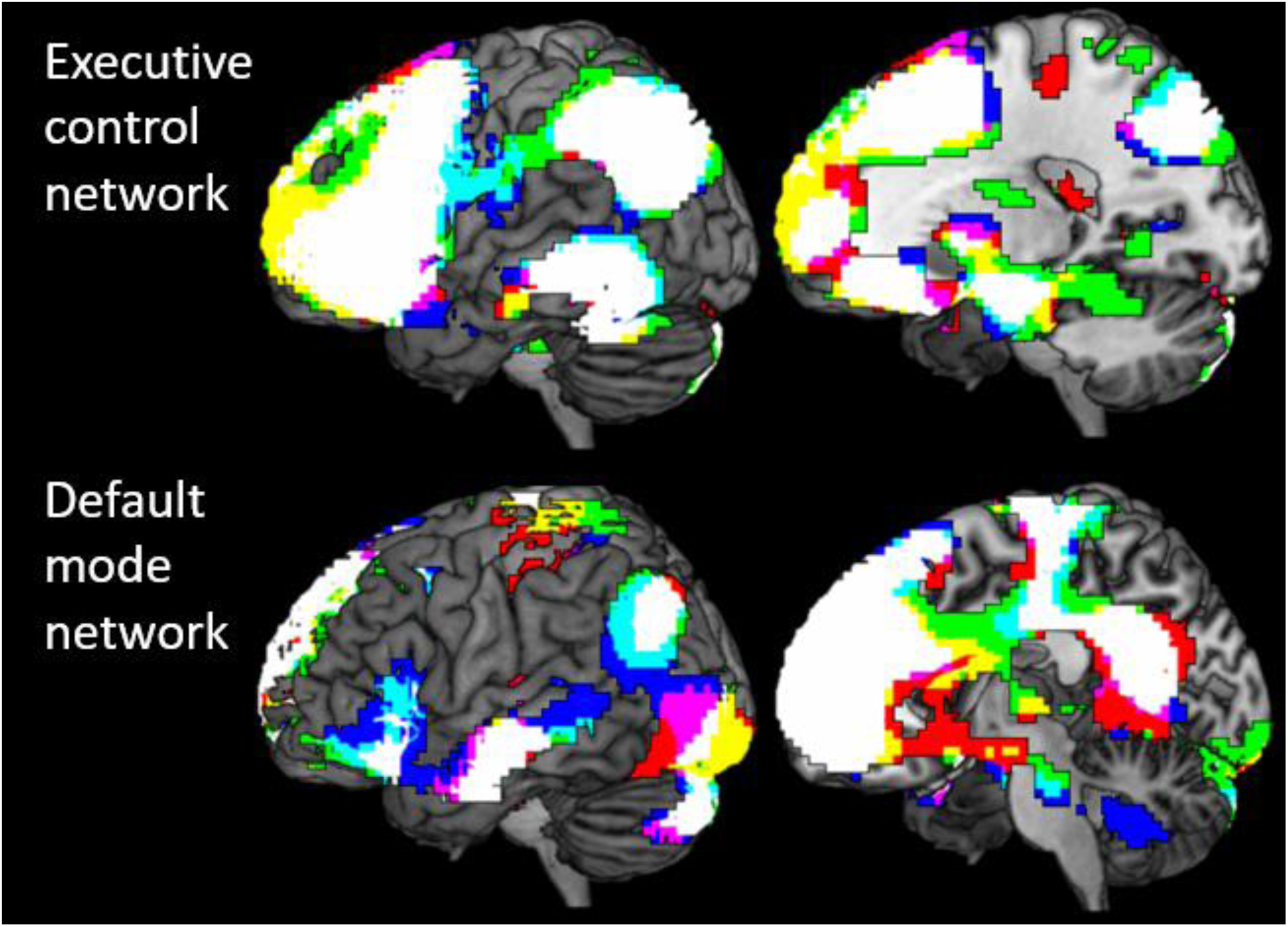
The executive network and default mode network for each task: numbers (red), picture (blue), and sentences (green). The overlap between networks is shown in white.

**Supplementary Figure 2.**
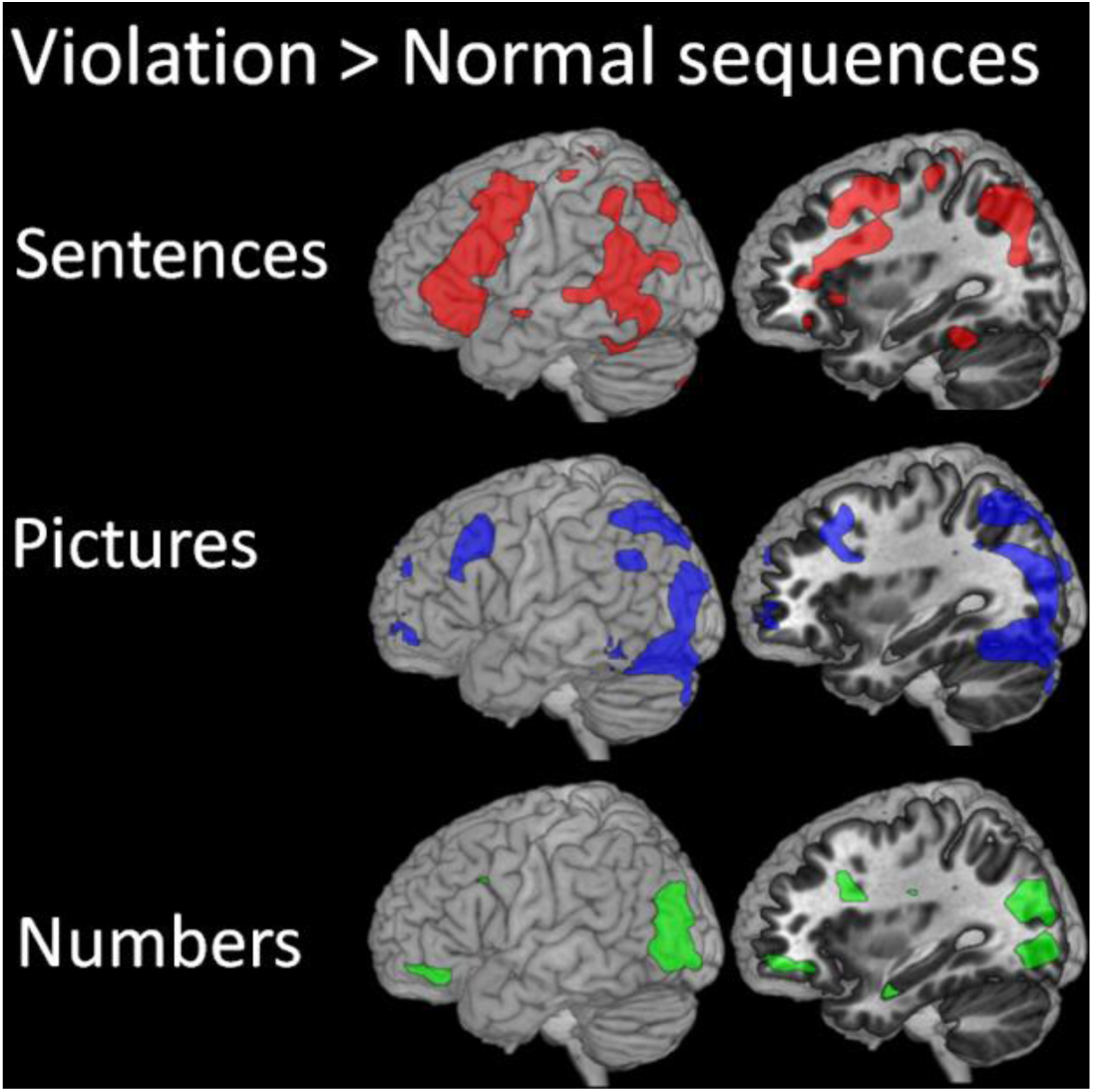
The results from the GLM analysis for the Violation > normal sequence contrast for each task (uncorrected, p < .001 for visual purposes).

**Supplementary Table 1.** The locations of the activation peaks from the GLM analysis.

|  | cluster<br>size | T | x | y | z | Location |
| --- | --- | --- | --- | --- | --- | --- |
| Sentences: |  |  |  |  |  |  |
| Violation > normal | 920 | 9.15 | -42 | 21 | 21 | Inferior frontal Gyrus, BA44 |
|  |  | 9.13 | -48 | 18 | 15 |  |
|  |  | 6.59 | -39 | 3 | 33 |  |
|  | 505 | 7.58 | -51 | -54 | 15 | Angular gyrus |
|  |  | 5.19 | -33 | -42 | -21 | Temporal fusiform |
|  |  | 5.04 | -63 | -51 | 3 | Middle temporal gyrus |
|  | 184 | 5.73 | -30 | -69 | 45 | Angular gyrus, PGa |
|  |  | 5.73 | -27 | -75 | 39 | Superior parietal lobule |
|  | 66 | 5.37 | -12 | -33 | 69 | Postcentral gyrus |
|  | 191 | 5.16 | 48 | -48 | 9 | Angular gyrus, PGa |
|  |  | 5.10 | 51 | -30 | 0 | Superior temporal gyrus |
|  |  | 4.52 | 57 | -42 | 6 | Middle temporal gyrus |
|  | 122 | 4.97 | 48 | 3 | 30 | Precentral gyrus |
|  |  | 4.61 | 48 | 15 | 21 | Inferior frontal Gyrus, BA44 |
|  | 39 | 4.71 | -9 | 18 | 45 | Superior frontal gyrus |
|  | 25 | 4.40 | -36 | -69 | 18 | Angular gyrus, PGp |
|  | 24 | 4.38 | 33 | -66 | 45 | Superior parietal lobule |
| Numbers: |  |  |  |  |  |  |
| Violation > normal | 84 | 5.08 | -39 | -75 | -3 | Lateral occipital cortex |
|  |  | 3.77 | -27 | -90 | -6 | Occipital pole |
|  | 32 | 4.76 | -12 | -36 | -36 | Cerebellum |
|  |  | 4.06 | 0 | -33 | -39 |  |
|  |  | 4.00 | 12 | -30 | -36 |  |
|  | 81 | 4.42 | -36 | -78 | 24 | Angular gyrus, PGp |
|  |  | 4.08 | -24 | -78 | 24 |  |
| Pictures: |  |  |  |  |  |  |
| Violation > normal | 2709 | 7.09 | 24 | -57 | -12 | Fusiform gyrus |
|  |  | 6.79 | 33 | -75 | -15 |  |
|  |  | 6.69 | 18 | -78 | -18 |  |
|  |  | 6.65 | -24 | -81 | -15 | Fusiform gyrus |
|  |  | 6.56 | -36 | -84 | -18 | Lateral occipital cortex |
|  |  | 5.95 | -9 | -81 | -6 | Lingual gyrus |
|  |  | 6.40 | -18 | -72 | 48 | Superior parietal lobule, 7P |
|  |  | 5.85 | -27 | -81 | 18 | Angular gyrus, PGp |
|  | 170 | 6.72 | 3 | -42 | 15 | Posterior cingulate |
|  |  | 5.49 | -3 | -24 | 27 | Posterior cingulate |
|  | 111 | 5.93 | 45 | 24 | 39 | Middle frontal gyrus |
|  | 65 | 5.83 | 51 | -51 | 33 | Angular gyrus, PGa |
|  |  | 3.74 | 51 | -51 | 45 |  |
|  | 81 | 4.86 | -39 | 18 | 45 | Inferior frontal gyrus, BA44 |
|  | 19 | 4.44 | -51 | -57 | 30 | Angular gyrus, PGa |
|  | 23 | 4.19 | 33 | 60 | -6 | Frontal pole |
|  |  | 3.68 | 45 | 51 | -6 |  |
| Violation effect: Sentences > numbers & pictures | 75 | 4.33 | -36 | -3 | 33 | Precentral gyrus |
|  |  | 4.26 | -39 | 0 | 51 |  |
|  | 41 | 4.28 | -39 | 18 | 21 | Inferior frontal gyrus, BA44 |
|  | 45 | 4.15 | -54 | -51 | 21 | Angular gyrus, PGa |
|  | 30 | 3.86 | 54 | -30 | 0 | Superior temporal gyrus |
|  |  | 3.84 | 54 | -36 | 6 |  |
| Violation effect: Pictures > sentences & numbers | 102 | 4.70 | 30 | -63 | -12 | Fusiform gyrus |
|  |  | 4.60 | 30 | -78 | -9 |  |
|  |  | 3.82 | 15 | -81 | -12 |  |
|  | 44 | 4.40 | -18 | -81 | -12 | Fusiform gyrus |
|  | 51 | 3.89 | 12 | -90 | 18 | Visual cortex |

